# Resting Eyes Closed Beta-Phase High Gamma-Amplitude Coupling Deficits in Children with Attention Deficit Hyperactivity Disorder

**DOI:** 10.1101/598003

**Authors:** David Ibanez-Soria, Eleni Kroupi, Andrés Rojas, Marta Castellano, Jacobo Picardo, Gloria García-Banda, Belen Saez, Mateu Servera, Giulio Ruffini, Aureli Soria-Frisch

**Author notes:** **Corresponding Author:** David Ibáñez Soria, Starlab Barcelona S.L., Av. del Tibidabo, 47 bis, 08035 Barcelona, Spain.

## Abstract

**Objective:** Attention-deficit hyperactivity disorder (ADHD) is the neurobehavioral disorder with the largest prevalence rate in childhood. ADHD is generally assessed based on physical examination of the child and interviews, and therefore prone to subjectivity. This fact may lead to a high risk of mis- and over-diagnosis, a problem that can be addressed through the use of objective markers.

**Methods:** Here we propose to use phase-amplitude coupling as a digital biomarker in ADHD. We investigated the hypothesis that coupling between the phase of slow brain rhythms and the amplitude of fast rhythms is altered in the ADHD population. We tested this hypothesis measuring phase-amplitude coupling (PAC) in the 4 to 200Hz range in electroencephalographic (EEG) signals recorded in the central-frontal area in children during eyes closed resting state.

**Results:** Using automatic clustering, we observed statistically significant beta-gamma PAC deficits in the ADHD population in the frontal-left hemisphere.

**Conclusions:** These findings suggests alterations in the beta-gamma coupling in the ADHD population. We discuss the hypothesis that these alterations may be indicators of working memory and attention deficits.

**Significance:** The study of the coupling between the different brain rhythms can potentially contribute to the understanding and clinical diagnosis of ADHD.

## 1. Introduction

Attention-Deficit Hyperactivity Disorder (ADHD) is one of the most common neurodevelopmental disorders of childhood with an estimated prevalence rate between 5-7% (Willcutt 2012). ADHD symptoms have a severe impact on the lives of children, negatively influencing their academic performance and their social and family relationships. These symptoms are generally grouped as inattention, hyperactivity and impulsivity, whose appearance varies from patient to patient. Based on these symptoms, three subtypes of ADHD are diagnosed: 1) Predominantly hyperactive/impulsive (ADHD-H), 2) Predominantly inattentive (ADHD-I) and Combined subtype (ADHD-C) (A.P. Association 2013).

ADHD diagnosis is in general assessed based on physical examination of the child and interviews with the child, family and teachers (A.P. Association 2013, A. Diagnostic 2000, W.H. Organiztion 2004). As pointed out by the scientific community, ADHD assessment is subjective and heavily biased towards personal practice and experience, which leads to an intrinsic risk of mis- and over-diagnosis (Merten 2017, Crippa 2017). Thus, ADHD research is currently focused on the development of markers that support accurate clinical diagnosis and reduce the risk of over-diagnosis.

Electroencephalogram (EEG) has been used for decades for the characterization of abnormal neural activity in ADHD children (Lenartowicz 2014). Classical EEG analysis focuses on the study of differences in brain rhythms in the ADHD population compared to age-matched control groups. Many studies report increased theta band power (Snyder 2015, Ogrim 2011) and decreased power in the beta band (Buyck 2014, Clarke 2007) in the frontocentral brain region in the ADHD population. The theta/beta power ratio (TBR) that combines these two metrics has long been used as an ADHD biomarker (Ogrim 2011). Neuropsychiatric EEG-Based ADHD Assessment Aid (NEBA®), which combines the analysis of TBR with clinician’s evaluation, is an approved methodology by the US Food and Drug Administration (FDA) to support the diagnosis of ADHD [12]. However, many studies have reported insufficient accuracy for TBR and theta power in distinguishing children with ADHD from their control group (Buyck 2014, Liechti 2013). The discovery of robust, accurate ADHD biomarkers, thus, remains an open research issue.

EEG measures the combined electrical activity generated by large population of neurons that fire in synchronization (Silva 2013). This electrical activity has been traditionally characterized by its oscillatory nature, and is altered in several neurodevelopmental disorders including ADHD (Ibáñez 2018). EEG analysis traditionally studies the stationary activity of brain oscillations by segmenting the signal coming from an electrode into short-time epochs that are considered to be pseudo-stationary. The power at specific frequency rhythms within these epochs is then estimated (Cohen 1977) and subsequently averaged (Welch 1967). Power within specific rhythms has proved to be correlated with different physiological, cognitive or pathological conditions (Tatum 2014). However, the nature of the brain (Skarda 1987) advocates for the use of feature extraction techniques capable of detecting richer signal dynamics. Cross Frequency Coupling (CFC) analysis is capable of distinguishing non-stationary relationships between oscillations at different frequencies. Among them Phase amplitude coupling (PAC) characterizes the non-linear relationship between the phase and the amplitudes across frequencies and has been shown to be useful in different neurological and mental disorders (De Hemptine 2013, Swann 2015, Ahmadi 2019, Berman 2015, Hirano 2018).

PAC has gained enormous popularity in the recent years as a means to understand how the amplitude and the phase of distinct oscillations regulate dynamic communication within the brain (Seymour 2017). CFC of neural oscillations is not entirely understood and is currently an active field of research in neuroscience. Several studies have demonstrated coupling between the phase of low frequencies and the amplitude of fast rhythms (Canolty 2010), and that such coupling is altered in several neural diseases. PAC has been studied in neurodegenerative diseases such as Parkinson’s (De Hemptine 2013, Swann 2015), where elevated synchronization between the beta-phase and the broadband-gamma amplitude has been observed. In (Ahamadi 2019), a computer aided diagnosis system for early multiple sclerosis was devised using phase-amplitude coupling as biomarker. PAC has also been analyzed in neurodevelopmental diseases such as Autism Spectrum Disorders (Berman 2015), reporting abnormal alpha-gamma coupling during resting-state EEG, or schizophrenia (Hirano 2018). PAC has also been used to discriminate between healthy and pathological populations in ADHD. For instance, in ADHD, a reduction of the theta-gamma coupling during attention demanding tasks (Kim 2016) has been reported, and in (Kim 2015) the ADHD group exhibited a decrease in theta-gamma coupling in frontal, temporal and occipital areas during resting-state EEG (eyes open). The suitability to use PAC in ADHD characterization comes from the fact that coupling between oscillations has been proved to be a brain mechanism in attention (Mento 2018, Szczepanski 2014), sensory processing (Seymour 2017), or working memory (Chacko 2018).

However, to the best of our knowledge, PAC has not been used for the investigation of EEG eyes-closed resting-state of ADHD patients. Here, we aim to test whether the decrease in coupling observed in eyes open and attention-demanding tasks extends to eyes-closed EEG. We apply a non-parametric clustering method to identify significant PAC clusters that account for the multiple-comparison problem. This methodology allows to detect statistical significant regions without prior assumptions on the bands of interest. This approach was used in a recent paper (Chacko 2018), where the authors estimated PAC and identified significant clusters using ECoG in an epileptic population during a visual cognitive task. The PAC algorithm differs from the one used in this study: while in (Chacko 2018) the coupling modulation index is calculated following the entropy based approach proposed by Tort (Tort 2010), in this study we followed the modulation index methodology introduced by Canolty (Canolty 2006) as explained in section 2.3. Thus, our study differs substantially from the current state-of-the art.

Given the limited number of cross-frequency coupling studies in ADHD, and the potential of this feature for the objective characterization of this disorder, this work aims to deepen the study of phase amplitude coupling abnormalities in the ADHD population compared to an age-matched healthy control population during resting eyes closed. We have investigated coupling anomalies in brain rhythms in the 4 to 200 Hz range to include the whole EEG spectrum ranging from theta to high gamma. We demonstrate that ADHD children present statistically significant deficits in beta-phase high-gamma-amplitude coupling in the frontal cortex. This metric thus represents a potential neurophysiological marker for ADHD.

## 2. Materials and Methods

### 2.1 Demographics

In this study, 51 children (aged 7 to 11 years) participated in an EEG recording after obtaining signed parental informed consent, where a detailed explanation of the study was provided. Participants were divided in two groups: clinical diagnosed ADHD group and healthy controls.

Children diagnosed with ADHD were recruited from specialized pediatric disorders clinical units in Palma de Mallorca, Spain. The inclusion criteria in the ADHD group were the following: (1) being clinically diagnosed with ADHD by a specialist using the DSM-IV criteria, (2) not having comorbidity problems of mental retardation, autism, bipolar or psychotic disorders, history of epileptic seizures or any other relevant medical disorder, and (3) refrain from ADHD medication 24-hours prior the EEG assessment. The ADHD group included 21 children: 12 with combined ADHD subtype (11 males and 1 female) and 9 with inattentive subtype (3 males and 6 females).

Healthy controls were selected from school age-matched standard classrooms. Inclusion criteria for this group were: (1) not having any psychopathology diagnosis, mental retardation or learning disorders, (2) not showing behavioral problems or learning difficulties in class (as asserted by their tutors) (3) not having major family problems that could interfere with their participation in the study. The control sample consisted of 30 children. The demographic characteristics of the experimental groups are summarized in Table 1.

**Table 1:** Experimental Population Demographic Information

|  | **ADHD (n=21)** | **Control (n=30)** |
| --- | --- | --- |
| **Age in Years** | MD 9.6 (SD 1.4) | MD 9.3 (SD 1.5) |
| **Male/Female** | 14/7 | 13/17 |
| **ADHD C/I** | 12/9 | - |
| Medication<br>(Yes/No) | 6/15 | - |

### 2.2 Experimental Procedure

Experimental recordings followed a 3-minutes resting state eyes closed protocol: participants were comfortably seated in a quiet room and were instructed to relax, keep their movements to a minimum and close the eyes during the EEG assessment. EEG data was recorded using a Neuroelectrics Enobio® recording system at a sampling rate of 500 S/s. The electrical reference (CMS/DRL) was placed in the right mastoid. As most abnormalities in the ADHD population were obtained in the fronto-central regions (Brooker 2016), we measured the brain activity of the participants in C3, Cz, C4, F3, Fz and F4 using Ag/AgCl electrodes according to the 10/10 EEG standard positioning system (Jurcak 2007).

**Figure 1.**
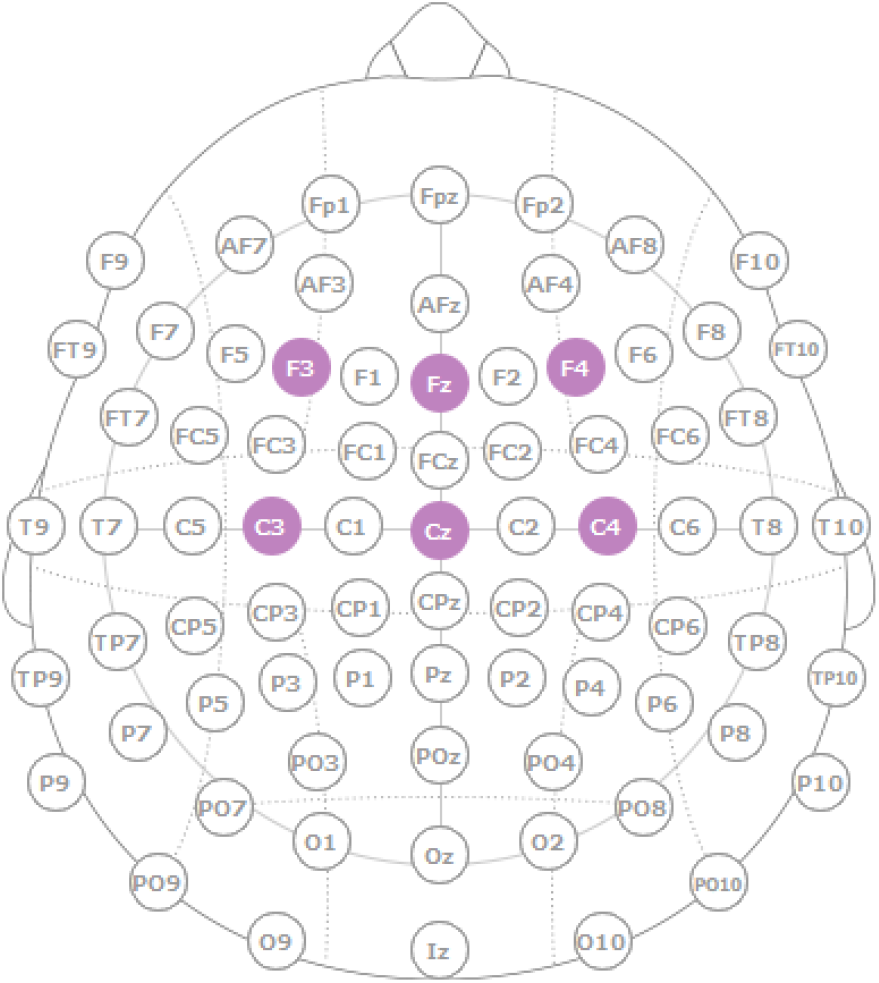
A graphical representation of electrode location.

### 2.3 Signal Processing and PAC Calculation

Phase amplitude coupling is computed following the modulation index (MI) approach proposed by Canolty et al. (Canolty 2006) and the implementation of Onslow et. al (Onslow 2011), and calculated for the frequencies ranging from 4 Hz to 200 Hz in 4 Hz steps.

First the signals from the 6 electrodes (C3, Cz, C4, F3, Fz, F4) are split into 30-second epochs with 50% overlap. To compute the coupling between the amplitude at the frequency band *a* and the phase at frequency band *p*, raw epochs of 30 sec, denoted as *e*(*t*), are first analyzed in the time-frequency domain through a wavelet convolution. This operation can be applied to bandpass filter the signals in the bands of interest (Cohen 2014). We apply a convolution with complex Morlet wavelets of width 7, which has been shown to overcome Infinite Impulse Response (IIR) filters in capturing the necessary temporal dynamics for the coupling characterization (Cohen 2014). The instantaneous amplitude *A_a_*(*t*) and phase *φ_p_*(*t*) are subsequently calculated as the absolute value and the phase angle of the complex-valued signal resulting from the filtering stage. Then the composite signal *z*(*t*) = *A_a_*(*t*) *e*^*jφ_p_*(*t*)^ is constructed.

If the probability density function of *z*(*t*), represented in the complex plane, is not radially symmetric (i.e., the mean of the *z*(*t*) is non-zero), then either *A_a_*(*t*) and *P_p_*(*t*) share mutual information or the distribution of *φ_p_*(*t*) is non uniform (Brooker 2016). Therefore, if the average of *z*(*t*) is non-zero and we assume that *φ_p_*(*t*) follows a uniform distribution, then there is coupling between *A_a_*(*t*) and *φ_p_*(*t*) (Jurcak 2007). This coupling is described by the modulation index, defined as the absolute value of the average *z*(*t*). Figure 2 displays the phase amplitude coupling calculation pipeline.

**Figure 2.**
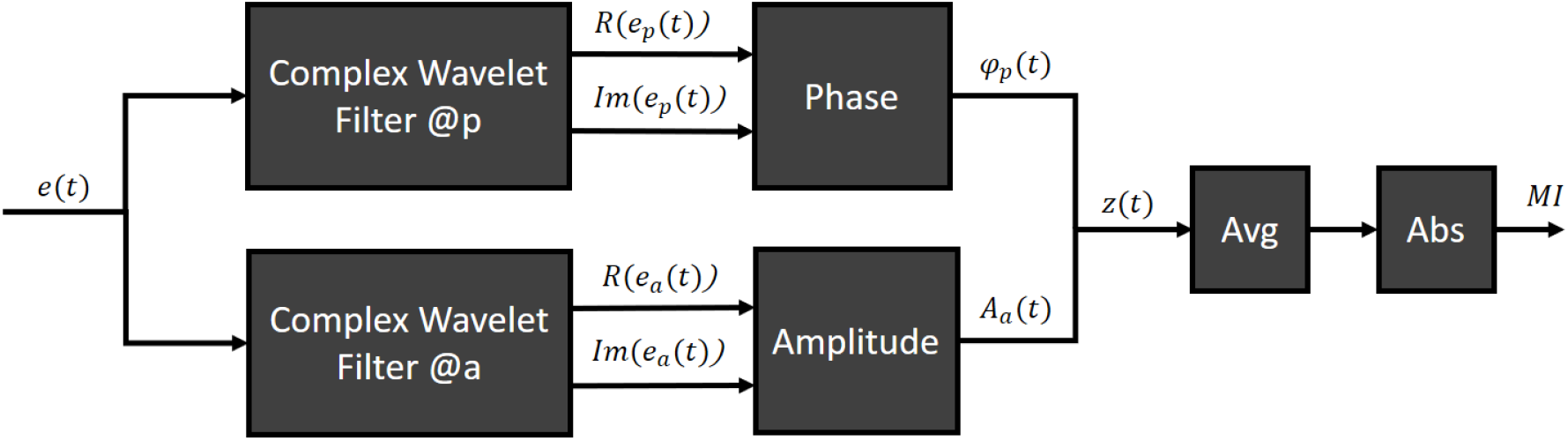
Phase amplitude coupling calculation pipeline

In order to remove the influence of the non-uniformity of *φ_p_*(*t*), shuffled datasets are used to conduct a significance analysis of the PAC values found (Onslow 2011). In particular, *A_a_*(*t*) is shuffled dividing the data into 1000 randomly selected sections that are then randomly rearranged to construct the shuffled signal. This way, we remove the temporal relationship of the amplitude values, whereas we preserve amplitude-related characteristics, such as mean, variance, and power spectrum. Following (Onslow 2011), 50 shuffled data sets are used with the significance value threshold at 0.05. Thus, PAC values lying at the 5% of the values distribution generated by the shuffled datasets are considered significant. Non-significant PAC values are set to zero. The final PAC matrices are therefore generated including the MI values where significant, and zero otherwise. These matrices are calculated at electrode level for each subject and epoch, and the final averaged PAC values over subjects and electrodes are computed at all artefact free epochs. Epochs are marked as noisy if they contain samples larger than 75uV in the 4 to 200Hz interval after power line noise removal. The final PAC matrices are subject to further non-parametric cluster analysis correcting for false positives, as indicated in the following section.

### 2.4 Non parametric Statistical Analysis

In this study we have used a non-parametric statistical test based on clustering, proposed by Maris and Oostenveld (Maris 2007), that addresses the multiple-comparison problem. The proposed method has been shown to perform well on MEG and EEG data. Thus, following (Maris 2007): 1) we draw random partitions of the data (matrix in the form of phase x amplitude PAC values) without replacement, creating new data matrices of the same size as the initial one; 2) we estimate the test statistic we are interested in, both on the initial data set as well as on the random partitions; and 3) we estimate the Monte-Carlo approximation of the p-value by accounting of the difference between the test statistic of the initial matrix and the test statistics of the random partitions. In this case we used the student t-test statistic (we computed an estimate of means and supposed unknown standard deviations for the distributions), with 1000 random partitions and a two-sided test.

In order to deal with the multiple-comparison problem, following (Maris 2007) we:

1. estimate one test statistic value for each PAC pair within each electrode,
2. selected all test statistic values above the 97.5% percentile to represent the most significant values,
3. clustered the test statistic samples in terms of adjacency (phase and frequency adjacency),
4. calculate the cluster-based statistics by taking the sum of the values within a cluster,
5. take the largest absolute value across all clusters to be the original test-statistic, and
6. apply the permutation test previously described for each PAC cluster to give us a final p-value for each cluster.

The contribution of the method to the non-Gaussian properties of the data, as well as to the multiple-comparison problem is further elaborated in Section 4.

This procedure is graphically presented in Figure 3. The matrices are generated only for visualization purposes. The actual results are presented in the following section. Since we only had six electrodes, we applied this procedure for each electrode.

**Figure 3.**
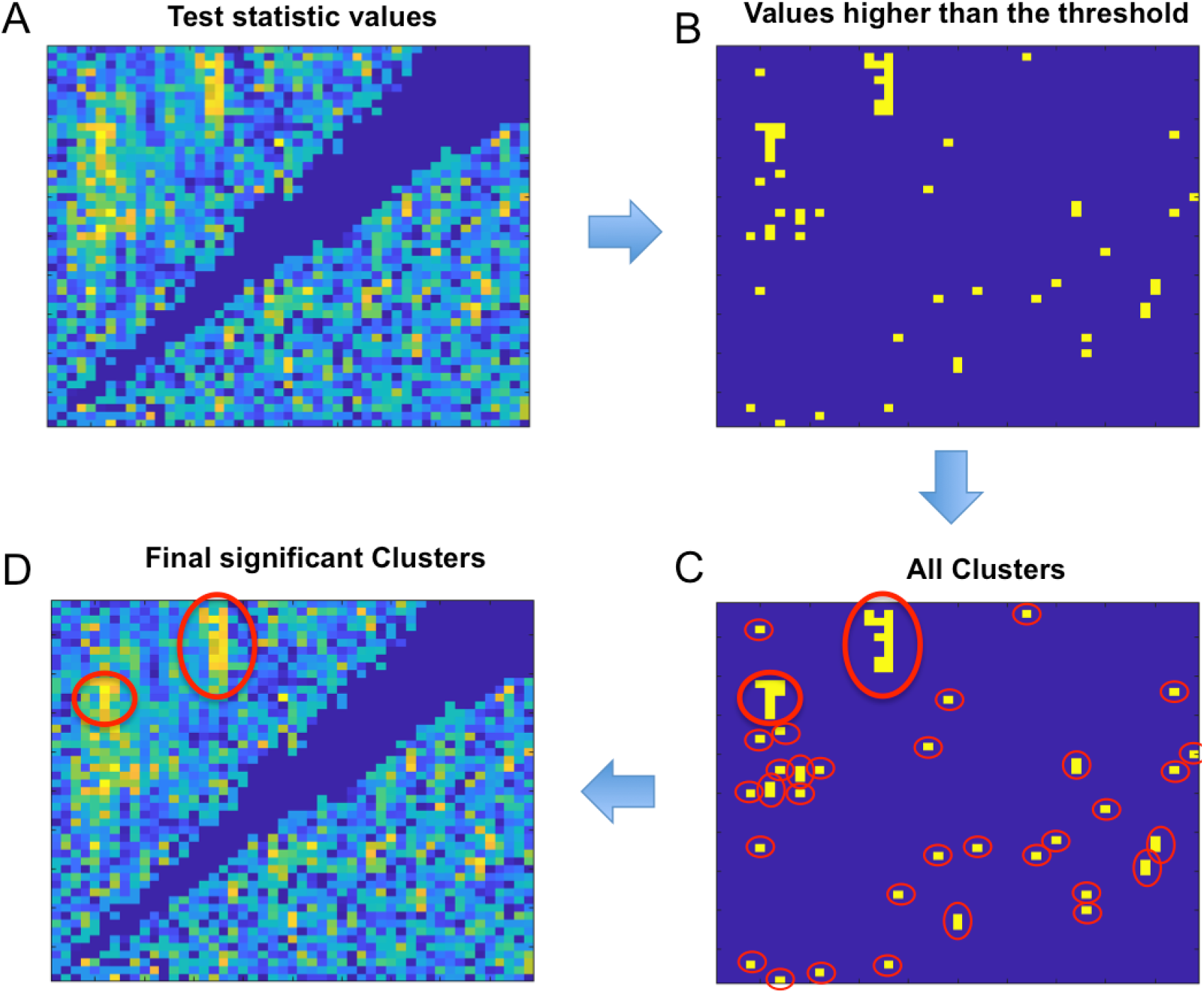
A graphical example of the non-parametric clustering procedure. Figure 3A displays the test statistic values for each PAC pair of an electrode. Figure 3B displays the test statistic values that are higher than the 97.5% percentile. Figure 3C presents all the adjacent clusters of the test statistic values. Figure 3D presents the final significant clusters.

## 3. Results

Relationships between phase and amplitude EEG rhythms have been computed at each electrode according to the previously presented cluster-based permutation test. This allows to investigate which brain regions are mainly responsible for changes in the ADHD population compared to healthy controls. Statistically significant differences between the ADHD and the control population were found in the electrode F3 (i.e., frontal left hemisphere). Figure 4 shows the grand average phase amplitude coupling in the 4 to 200Hz frequency range for the ADHD (left) and control (right) populations at F3. The region with statistical significance between groups is marked by a red contour.

**Figure 4.**
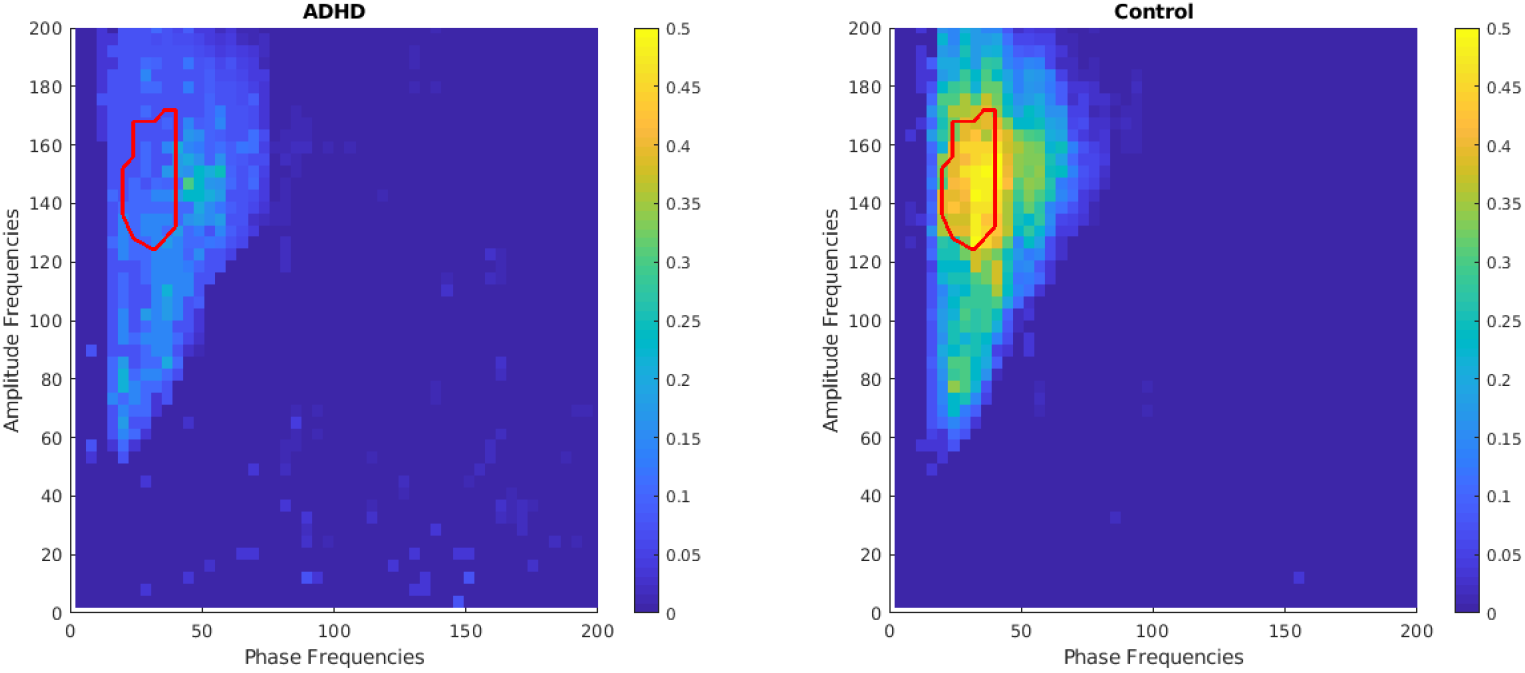
Grand average phase amplitude coupling of ADHD and Control populations at F3. The red contour displays the significant region as estimated with the non-parametric clustering algorithm.

Figure 5 shows the grand average phase amplitude coupling in intervals of interest, namely in the phase 4 to 50 Hz and the amplitude 100 to 180 Hz frequency ranges. We can observe that differences between groups are found in the beta phase (20-40 Hz) and high-gamma amplitude (128-174 Hz) coupling, with the ADHD showing a decrease in synchronization (p<0.05).

**Figure 5.**
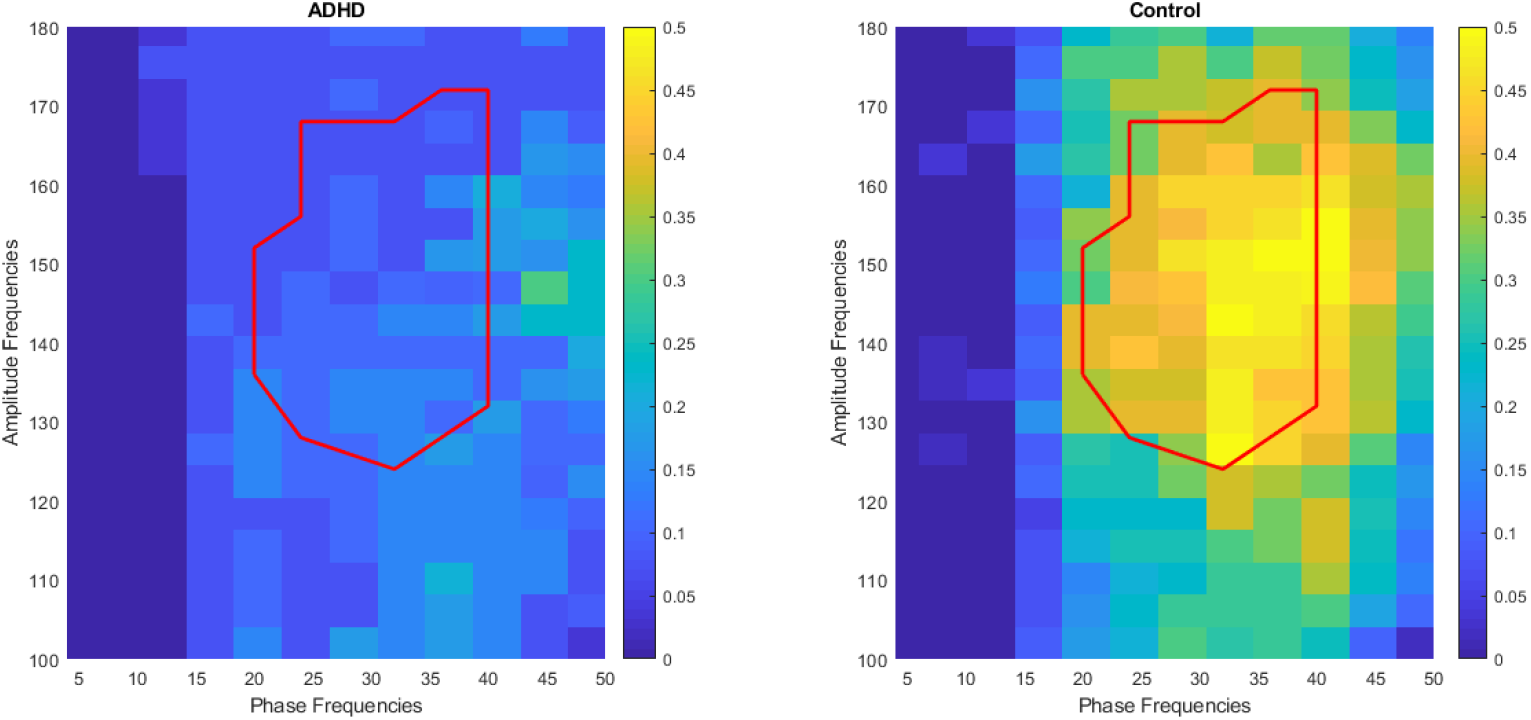
Grand average phase amplitude coupling of ADHD and Control populations at F3. The red contour displays the significant region as estimated with the non-parametric clustering algorithm.

We have calculated the average PAC of the area of interest obtained for F3 at every electrode. Figure 6 shows the grand average of the PAC feature along with its standard error of the mean over the ADHD and the control populations, and Figure 7 displays its topographical distribution. We can observe from both figures that although the beta-phase high-gamma-amplitude coupling deficit of the ADHD population was statistically significant only in F3, there is a trend of lower coupling in the ADHD population in the entire frontocentral region, although not significant (p>0.05).

**Figure 6.**
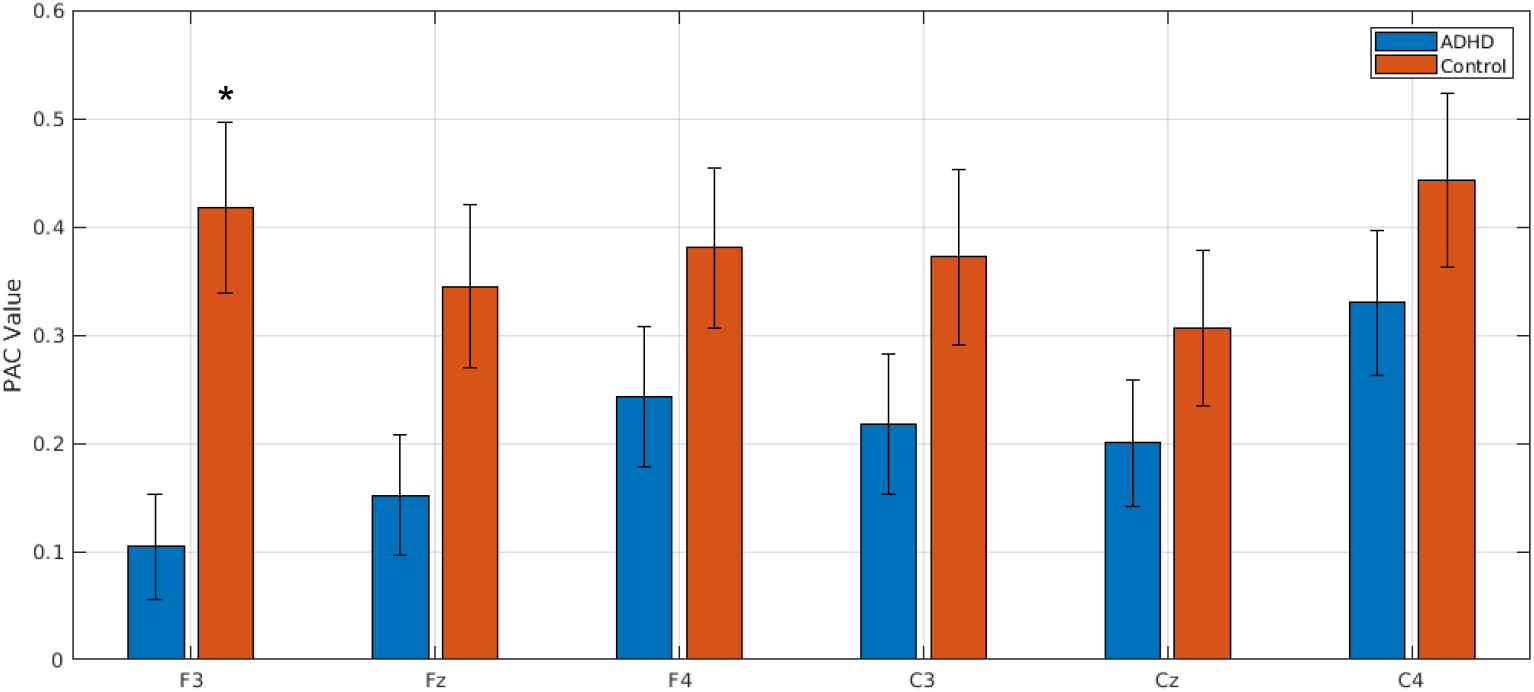
Beta-phase high-gamma-amplitude coupling ADHD and control population. The bar plots represent the grand average and the whiskers extend to the standard error of the mean.

**Figure 7.**
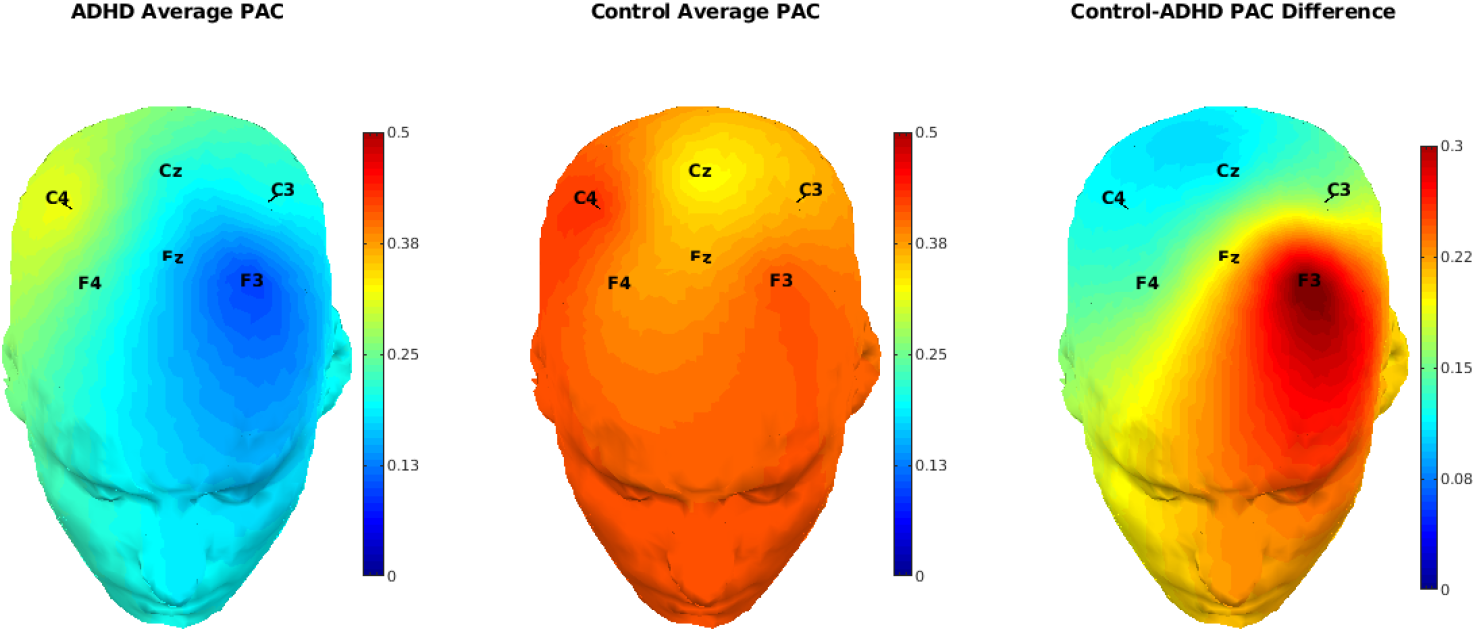
Topographical distribution of beta high-gamma PAC: Left) ADHD, Middle) Control Group, Right) Difference between ADHD and Control group.

## 4. Discussion

Given ADHD over-diagnosis (Bruchmuller 2012) and medication side effects (Graham 2011), it is important to find reliable physiological biomarkers that may help to improve ADHD diagnosis. In this work we have further investigated the use of PAC as a digital biomarker in ADHD, based on neurophysiological data. This is the first time to the best of the authors’ knowledge that PAC is studied in ADHD children during resting state and eyes closed. We have demonstrated overall beta-phase gamma-amplitude coupling deficits in frontal-central areas. These deficits proved to be statistically significant in the frontal-left hemisphere (channel F3).

Neurons do not work in isolation (Hebb 2005), and they rather tend to work in synchrony integrating widespread neural assemblies in a coordinated manner. The study of human interdependent neural oscillations has demonstrated a coupling between the phase of low frequencies and the amplitude of fast rhythms. According to (Buzsaki 2009), high-frequency oscillations may reflect local oscillatory patterns in small populations of neurons, whereas low-frequency activity may reflect large-scale functional connectivity among distal, large neuronal populations. PAC is capable of detecting complex interactions between these rhythms.

According to (Palva 2018), gamma oscillations may be related to bottom-up processes, whereas delta, theta, alpha and beta oscillations to top-down controlling functions. Thus, cross-frequency coupling properties among such oscillations can be indicators of coordination, integration, and regulation of separate neuronal assemblies (Staudigl 2012). Connections of fast and slow oscillatory networks, can, thus, show the integration of distributed cognitive and emotional functions, such as affective and sensory information, as well as attentional and executive functions (Palva 2018, Tort 2010). For instance, (Staudigl 2012) reported that the coupling of beta oscillations in the thalamus with gamma oscillations in the neocortex is related to performance in a recognition memory task, and (FitzGerald 2013) observed significant CFC between thalamus and neocortex for different brain regions, implying exchange of information flow between these two parts of the brain. (Tort 2010) also reports that alpha/beta oscillations of deep layers regulate gamma oscillations in superficial layers of the frontal cortex, and such regulations are related to the contents of working memory, as measured by local field potentials of monkeys. The frontal cortex is also related to cognitive functions, such as working memory, awareness, and attention, and provides top-down control of other brain regions (Palva 2018, Hefrich 2016, Riley 2016). Although our findings were extracted from resting-state EEG signals in humans and not under task performance, there still may be indications of working memory and attention deficits in the ADHD population, denoted in the frontal beta-gamma coupling reduction for this population.

Activity within a certain EEG band has been seen to correlate with different physiological or cognitive functions. Many definitions of EEG bands exist, but as of today there is no agreement in standard frequency ranges and band distributions are often considered arbitrary. Buzsaki, for example, defines in (Buzsaki 2009) this arbitrariness as “the straight-line country borders between the African nations drawn by the colonialists”. The definition of cross-frequency coupling regions of interest is in general assessed following one of these band definitions. In this study we used a cluster-based permutation test to detect statistically significant regions without prior assumptions on the bands. The proposed statistical method extracts the significant areas of interest by dealing with the multiple-comparison problem in an efficient way, which refers to the Type I errors introduced by carrying out multiple statistics. Concretely, the multiple comparison correction is attained with the 97.5% threshold (see Section 2.4), i.e. by selecting only the 2.5% extremely significant values, as well as by choosing the largest absolute value over all clusters to be the final test statistic and limiting this way the number of false positives. This improves the usual Bonferroni correction that often underperforms in correcting for the Type II errors. The proposed method also addresses the non-Gaussianity of the data issue, which is often the case for neuroscience data, through the proposed permutation test, that compares the test statistic of the data with the test statistics from randomly permutated data, without any prior assumption on the data distribution.

Using the same data-set, in a previous study we showed that the ADHD population presented exaggerated theta activity and that dynamical differences between eyes open and closed conditions were altered in the ADHD population in theta and beta (Ibáñez 2018 B). This work confirms the hypothesis that several brain patterns are altered in the ADHD, ranging from stationary rhythmic activity to complex dynamics and synchronization between channels. Authors plan to further explore cross-frequency coupling abnormalities in ADHD and ASD in larger data-sets.

## 5. Acknowledgements

We specially acknowledge the participants in this study. Funding has been partially provided by the European Commission through the STIPED project under the Horizon 2020 Program (H2020, Grant agreement number 731827). We thank Patricia Teruel and Marta García Pérez for their assistance in the EEG recordings.

